# H3K37me1 couples transcription with DNA replication origin selection

**DOI:** 10.64898/2026.06.22.733675

**Authors:** Ricardo de Arellano, Patrick Toolan-Kerr, Iria Domínguez-García, Diego Polanco-Alonso, Laura López-Hernández, Simone Sidoli, Néstor García-Rodríguez, Luca Pandolfini, Gonzalo Millán-Zambrano

**Affiliations:** Centro Andaluz de Biología Molecular y Medicina Regenerativa (CABIMER), CSIC-Universidad de Sevilla-Universidad Pablo de Olavide, Seville, Spain; Departamento de Genética, Facultad de Biología, Universidad de Sevilla, Seville, Spain; Department of Biochemistry, Albert Einstein College of Medicine, New York, New York, USA; Center for Human Technologies, Istituto Italiano di Tecnologia (IIT), Genova, Italy

## Abstract

DNA replication initiation in higher eukaryotes occurs at thousands of sites distributed throughout the genome and follows a defined temporal program. However, replication origins are not determined by a conserved DNA motif, and how specific genomic regions are selected for initiation remains poorly understood. Here, we identify a transcription-associated histone modification in human cells, mono-methylation of histone H3 lysine 37 (H3K37me1), and show that it regulates the spatial organization of replication initiation. H3K37me1 is enriched across actively transcribed gene bodies, and its depletion leads to a redistribution of replication origin activity toward intragenic regions. We show that H3K37me1 limits the association of the MCM2–7 replicative helicase with transcribed chromatin, thereby restricting intragenic origin usage. Consistently, loss of H3K37me1 increases both MCM2–7 occupancy and replication initiation across gene bodies. Collectively, our findings uncover a chromatin-based mechanism that couples transcription with DNA replication origin selection by limiting unscheduled origin firing within transcribed regions.

## Introduction

DNA replication initiates at numerous chromosomal sites termed replication origins, whose coordinated activation ensures faithful genome duplication. Replication origins are best characterized in *Saccharomyces cerevisiae*, where they are specified by a defined DNA consensus sequence predominantly located in intergenic regions.^1^ However, unlike in budding yeast, replication origins in higher eukaryotes are not determined by a conserved DNA motif, and how specific genomic regions are selected for initiation remains poorly understood.

Initiation of DNA replication involves two temporally separated steps.^2^ First, during late mitosis and G1 phase, the origin recognition complex (ORC), together with CDC6 and CDT1, loads the MCM2–7 replicative helicase onto chromatin to license potential replication origins.^3–5^ Upon entry into S phase, CDK and DDK activities promote the recruitment of limiting replication initiation factors that activate a subset of licensed sites to form functional replication forks.^6,7^ As a result, only a fraction of licensed origins initiate replication, thereby establishing a spatially organized DNA replication program across the genome.

Accumulating evidence indicates that local chromatin features, including specific histone post-translational modifications (PTMs), contribute to the control of replication origin activity.^8–10^ For example, methylation of histone H4 lysine 20 (H4K20) has emerged as an important regulator of replication origin function in metazoans.^11–13^ The bromo-adjacent homology (BAH) domain of ORC1 recognizes H4K20me2, and disruption of this interaction reduces ORC occupancy at replication origins.^12^ Similarly, H4K20me3 promotes recruitment of the ORC-associated protein ORCA (LRWD1), which stabilizes ORC binding and facilitates replication licensing at heterochromatic regions.^11^ However, while several histone PTMs have been linked to the activation of replication origins, comparatively less is known about the mechanisms that repress replication initiation across the genome.

In multicellular eukaryotes, the genome replicates in large chromosomal domains that follow a characteristic temporal order, establishing a replication timing (RT) program in which euchromatic regions replicate early and heterochromatic regions replicate late.^14^ Consistent with this organization, origin activity is closely associated with both chromatin accessibility and transcriptional output.^15,16^ Several studies have shown that, in unperturbed human cells, early replication initiation sites are frequently located in non-transcribed regions adjacent to the transcription start site (TSS) of actively transcribed genes.^15–18^ In contrast, replication initiation within active gene bodies is largely suppressed, thereby favoring the co-directional orientation of replication and transcription. In this regard, recent work has proposed that transcription redistributes MCM2–7 complexes to restrict early DNA replication initiation outside of active genes.^17–19^ Yet, the molecular mechanisms underlying this transcription-dependent relocation of MCM2–7 from gene bodies remain incompletely understood.

We previously showed that mono-methylation at lysine 37 (H3K37me1) regulates DNA replication in *Saccharomyces cerevisiae*.^20^ However, whether this modification controls replication origin selection in metazoan genomes remains unknown. Here, we report the presence of H3K37me1 in human cells, establish its association with transcription, and show that it regulates the spatial organization of DNA replication initiation. H3K37me1 limits MCM2–7 association with actively transcribed gene bodies, thereby restricting intragenic origin firing. Together, these findings uncover a chromatin-based mechanism that couples transcription with DNA replication origin selection.

## Results

### H3K37me1 marks actively transcribed gene bodies

Large-scale mass spectrometry studies have reported histone H3 peptides compatible with mono-methylation at lysine 37 (H3K37me1) in mammalian cells.^21–23^ However, its biological function has remained unknown. Because methylation of K36 and K37 produces identical mass shifts, unambiguous assignment of the modified residue requires fragment-ion patterns that resolve b/y ions between the two lysines. Using high-resolution MS/MS analysis of histones purified from hTERT-immortalized, untransformed human RPE-1 cells, we detected the H3 peptide encompassing residues 27–40 and manually validated fragment ions consistent with mono-methylation specifically at K37 (Fig. 1A), providing direct evidence for the presence of H3K37me1 in human cells.

**Figure 1.**
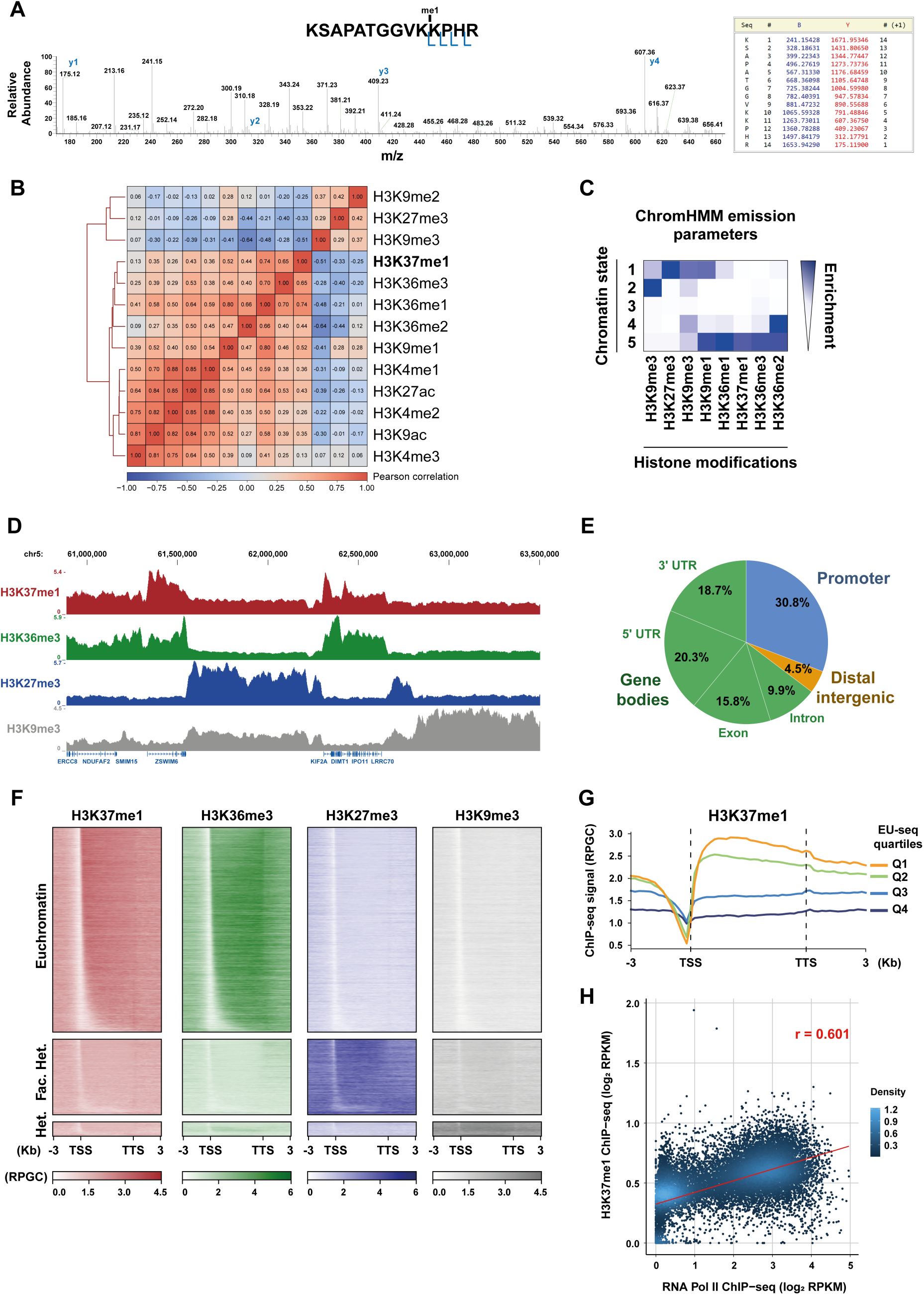
H3K37me1 marks actively transcribed gene bodies. **A.** MS/MS fragmentation spectrum of the H3 peptide (amino acids 27–40) with annotated y-ion fragments (y1–y4). The inset table lists theoretical b- and y-ion m/z values calculated for the peptide carrying mono-methylation at K37, which were used to match the observed fragment ions. **B.** Pearson correlation matrix for 13 histone PTMs. **C.** ChromHMM heatmap showing the emission probability of broad histone PTMs across five chromatin states. **D.** Genome browser snapshot of H3K37me1, H3K36me3, H3K27me3 and H3K9me3 ChIP-seq tracks. **E.** Distribution of H3K37me1 peaks across annotated genomic features. **F.** Heatmaps of H3K37me1, H3K36me3, H3K27me3, and H3K9me3 ChIP-seq signal across genes in euchromatin, facultative heterochromatin, and heterochromatin. Rows are sorted according to decreasing gene length. **G.** Metaplot of average H3K37me1 ChIP-seq signal along genes grouped into quartiles (Q1–Q4) by transcription level (EU-seq). **H.** Scatterplot of RNA Pol II (x-axis) versus H3K37me1 (y-axis) ChIP-seq signal at annotated genes.

To define H3K37me1 genomic distribution, we performed chromatin immunoprecipitation sequencing (ChIP–seq) experiments using our previously validated antibody.^20^ Importantly, the histone PTM landscape of RPE-1 cells has been extensively characterized,^24^ enabling direct comparison with a broad panel of euchromatic (H3K4me1/2/3, H3K36me1/2/3, H3K9ac, H3K27ac) and heterochromatic (H3K9me1/2/3, H3K27me3) modifications. Correlating H3K37me1 signal with these profiles revealed a close clustering with euchromatic marks (Fig. 1B), particularly those enriched across gene bodies, suggesting that H3K37me1 is preferentially associated with transcriptionally active chromatin. To further annotate the chromatin context of H3K37me1, we used ChromHMM^25^ and defined five chromatin states based on the combinatorial patterns of broad histone PTMs (H3K9me1/2/3, H3K27me3 and H3K36me1/2/3). Interestingly, H3K37me1 was assigned to the active chromatin state, with no representation in H3K27me3- or H3K9me3-dominated regions (Fig. 1C). Genome browser inspection of representative loci highlighted co-enrichment of H3K37me1 with H3K36me3 and its exclusion from repressive domains (Fig. 1D), reinforcing its euchromatic association.

Next, we performed peak calling analysis and found that H3K37me1 is almost exclusively detected at annotated genes (Fig. 1E), with minimal representation in distal intergenic regions. Consistent with this, we noticed that H3K37me1 is evenly distributed along euchromatic gene bodies (Fig. 1F), whereas no obvious enrichment is observed at enhancers (Fig. S1A). Metagene analyses further showed that H3K37me1 levels at gene bodies scale with transcriptional output measured by EU-seq^26^ (Fig. 1G), correlating positively with RNA polymerase II (RNA Pol II) occupancy^27^ (Fig. 1H) and chromatin accessibility as assessed by ATAC–seq^24^ (Fig. S1B). Altogether, these results indicate that H3K37me1 selectively marks actively transcribed gene bodies.

### H3.3K37R mutation selectively reduces H3K37me1 levels

A major obstacle to defining the biological function of individual histone PTMs is that histone-modifying enzymes often target multiple residues and frequently act on non-histone substrates.^28^ Consequently, definitive assignment of PTM-specific functions requires direct genetic manipulation of the modified residue.^29,30^ This approach has historically been difficult in mammalian cells because canonical histone genes are present in many copies. Recent work, however, has demonstrated that transcription-associated histone PTMs can be efficiently depleted by mutating the corresponding residue in the H3.3 histone variant, which is predominantly incorporated into actively transcribed genes through replication-independent deposition.^31,32^ Building on this principle, we used CRISPR/Cas9 to introduce a lysine-to-arginine substitution at position 37 of H3.3, a change that preserves the residue’s positive charge while preventing its methylation. Histone H3.3 is encoded by two genes, H3F3A and H3F3B, with the latter accounting for nearly 90% of total H3.3 expression in RPE-1 cells (Fig. S2A). Accordingly, we generated cells lacking H3F3A and carrying homozygous K37R mutations in H3F3B (H3F3A–/–; H3F3BK37R/K37R) and compared them with matched controls (H3F3A–/–; H3F3BWT/WT) (Fig. S2B). Correct editing of the H3F3B locus was confirmed by Sanger sequencing (Fig. S2C).

Next, we assessed the impact of the H3.3K37R mutation on chromatin-associated H3K37me1. Spike-in–normalized ChIP–seq (qChIP-seq) analysis revealed a notable depletion of H3K37me1 levels across euchromatic genes in the H3.3K37R mutant compared to the wild-type control (Fig. 2A), further validating the specificity of the antibody. This reduction extended uniformly along gene bodies (Fig. 2B) and was consistently observed across the entire gene set (Fig. 2C), indicating that the mutation results in a global rather than locus-specific loss of H3K37me1. Importantly, qChIP-seq profiling of H3.3 showed that neither its genomic distribution nor its overall levels were altered in mutant cells (Fig. 2D and S2D), arguing that the K37R substitution does not impair H3.3 incorporation into chromatin. Genome browser inspection further illustrated the selective loss of H3K37me1 at representative loci, with no detectable change in H3.3 occupancy (Fig. 2E). To exclude potential effects on the adjacent K36 residue, we profiled H3K36me1 and H3K36me3 by qChIP–seq. Both marks displayed virtually unchanged genomic distributions and signal intensities in H3.3K37R cells (Fig. 2F and S2E), suggesting that the K37R substitution does not perturb neighboring K36 methylation states. Similarly, histone mass spectrometry analysis showed no major alterations in global histone PTMs levels (Fig. 2G and S2F). Together, these results indicate that H3.3K37R selectively depletes H3K37me1 levels, validating this mutant as a precise tool to dissect the biological function of this modification.

**Figure 2.**
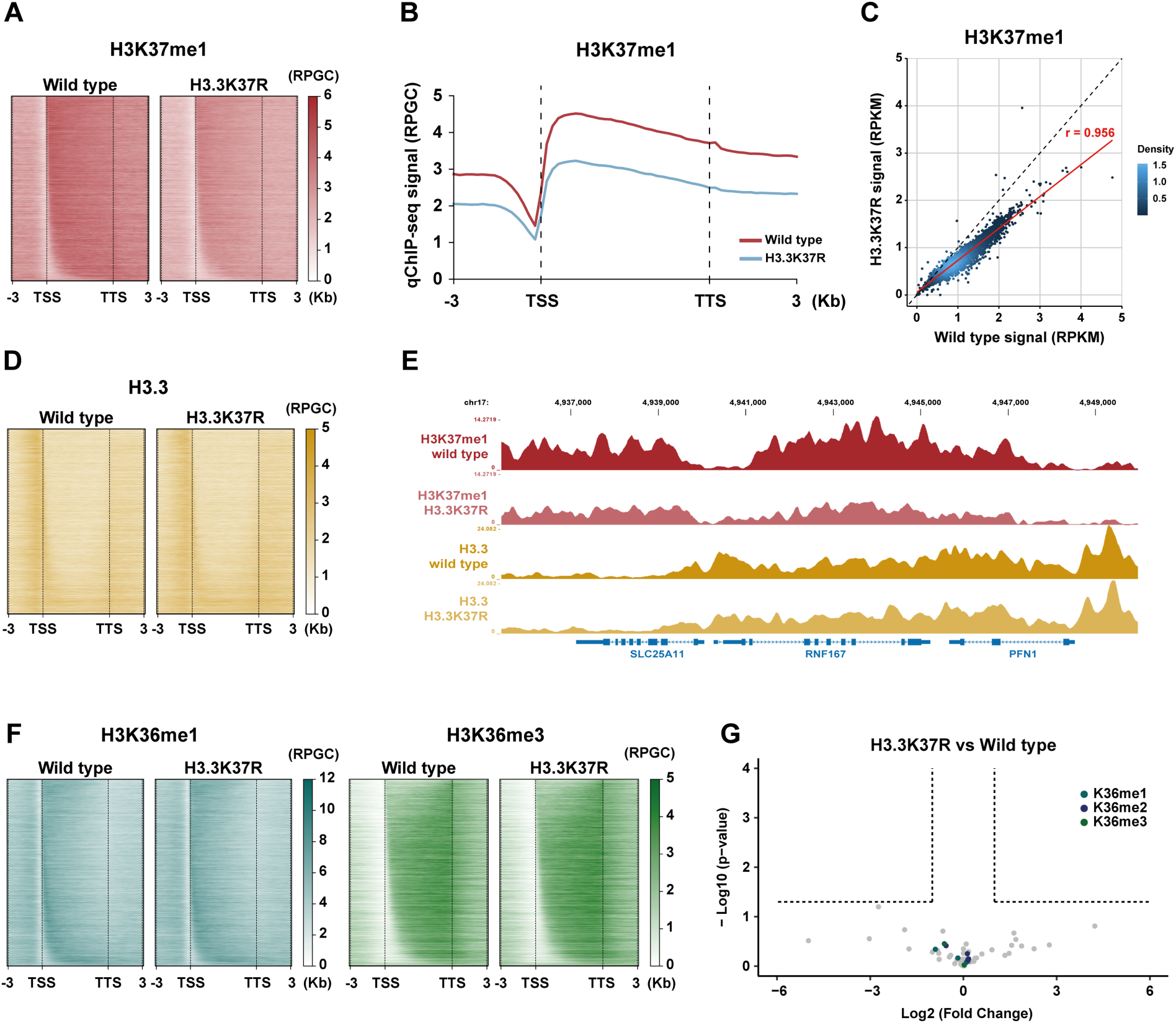
H3.3K37R mutation selectively reduces H3K37me1 levels. **A.** Heatmap of H3K37me1 qChIP-seq signal across euchromatic genes in wild-type and H3.3K37R mutant cells. Rows are sorted according to decreasing gene length. **B.** Metaplot of average H3K37me1 qChIP-seq signal across euchromatic genes in wild-type and H3.3K37R mutant cells. **C.** Scatterplot comparing H3K37me1 qChIP-seq signal across euchromatic genes in wild-type (x-axis) and H3.3K37R (y-axis) mutant cells. **D.** Metaplot of average H3.3 ChIP-seq signal across euchromatic genes in wild-type and H3.3K37R mutant cells. Rows are sorted according to decreasing gene length. **E.** Genome browser snapshot of H3K37me1 and H3.3 qChIP-seq tracks in wild-type and H3.3K37R mutant cells. **F.** Heatmap of H3K36me1 and H3K36me3 qChIP-seq signals across euchromatic genes in wild-type and H3.3K37R mutant cells. Rows are sorted according to decreasing gene length. **G.** Volcano plot displaying global changes in histone H3 post-translational modifications detected by mass spectrometry analysis in H3.3K37R versus wild-type cells. Two-sided Student’s t-test; n = 3 independent replicates; |log_2_ FC| > 1; p-value < 0.05.

### Loss of H3K37me1 does not globally alter transcription

Given the strong association of H3K37me1 with actively transcribed genes, we first examined whether this mark regulates transcriptional activity. To this end, we performed chromatin-associated RNA sequencing (chrRNA-seq) experiments.^33,34^ Global metagene and heatmap analyses revealed highly similar transcription profiles across gene bodies between wild-type and H3.3K37R mutant cells (Fig. 3A-B). Likewise, stratification by baseline transcriptional output showed comparable chrRNA-seq signal across all expression quartiles (Fig. S3A), indicating that loss of H3K37me1 does not result in detectable changes in transcriptional output. Consistently, only a limited number of transcripts displayed statistically significant expression changes (Fig. 3C), and gene ontology enrichment analysis identified categories unrelated to transcriptional regulation (Fig. 3D). Together, these results suggest that partial depletion of H3K37me1 does not broadly alter RNA Pol II–dependent transcription in RPE-1 cells.

**Figure 3.**
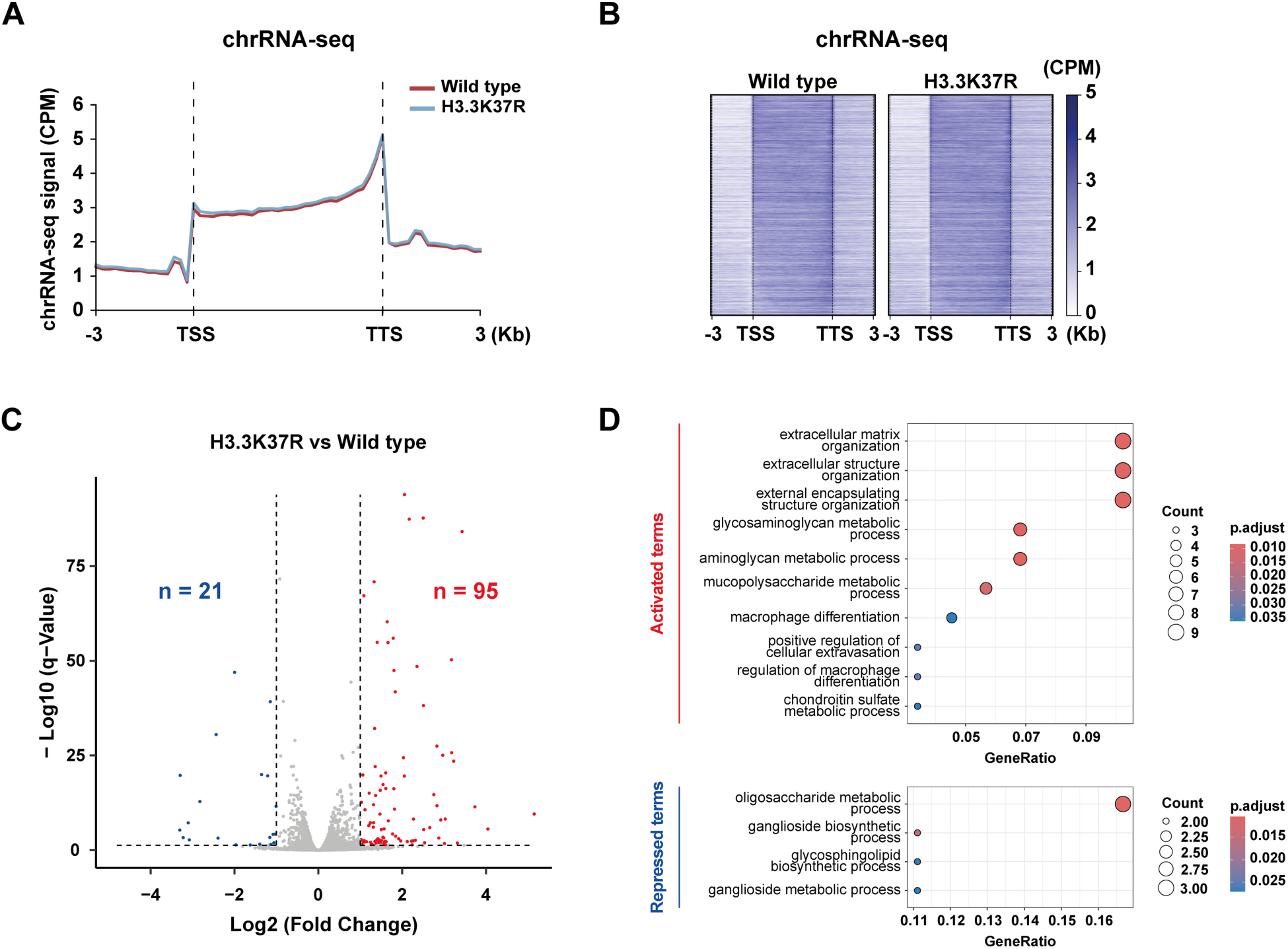
Loss of H3K37me1 does not globally alter transcription. **A.** Metaplot of average chrRNA-seq signal across euchromatic genes in wild-type and H3.3K37R mutant cells. **B.** Heatmap of chrRNA-seq signal across euchromatic genes in wild-type and H3.3K37R mutant cells. Rows are sorted according to gene length. **C.** Volcano plot showing differentially expressed genes in H3.3K37R mutant versus wild-type cells. Significantly up-regulated (red) and down-regulated (blue) genes are shown (|log_2_ FC| > 1, adjusted p < 0.05). **D.** Dotplot showing enriched biological processes among significantly up-regulated (red) and down-regulated (blue) gene sets.

### H3K37me1 restricts intragenic origin usage

In human cells, early DNA replication initiation sites are frequently located in non-transcribed regions adjacent to the TSS of actively transcribed genes, whereas replication initiation within gene bodies is generally suppressed. ^15–18^ We therefore asked whether transcription-associated H3K37me1 contributes to replication origin control in RPE-1 cells. To assess DNA replication initiation genome-wide, we performed EdU-seq-HU experiments.^35^ Wild-type and H3.3K37R mutant cells were synchronized in G1 using palbociclib and released into S phase in the presence of 5-ethynyl-2′-deoxyuridine (EdU) to label nascent DNA and hydroxyurea (HU), to limit fork progression (Fig. S4A).^36^ EdU-labelled DNA, which corresponds to early replication events, was isolated and subjected to high-throughput sequencing. Importantly, EdU-seq-HU profiles from wild-type cells closely matched previously published early replication initiation maps in RPE-1 cells (Fig. S4B and S4C),^17^ supporting the robustness of the experimental setup.

Peak calling analysis identified 6636 EdU-seq–HU peaks in wild-type cells (Fig. 4A), in line with previous reports.^17^ Notably, H3.3K37R cells showed a marked increase in the number of EdU-seq–HU peaks up to 9581, suggesting that loss of H3K37me1 enhances origin activation. To determine whether this increase reflects bona fide replication initiation events, we performed DNA fiber assays in wild-type and H3.3K37R mutant cells. Quantification of first-label origins —defined as IdU tracks flanked by CldU and used as a proxy for origin density—^37^ confirmed increased initiation events in H3.3K37R mutant cells (Fig. 4B). Fork progression rates were not reduced (Fig. 4C), indicating that the elevated origin density is not a secondary consequence of decreased fork speed. Moreover, because DNA fiber assays were performed in the absence of HU, these data argue against deregulated dormant origin activation as an explanation for the observed increased initiation frequency in H3.3K37R mutant cells. Importantly, these findings were further validated using a second independent H3.3K37R mutant clone (Fig. S4D-F). Altogether, the above results suggest that loss of H3K37me1 increases replication origin firing.

**Figure 4.**
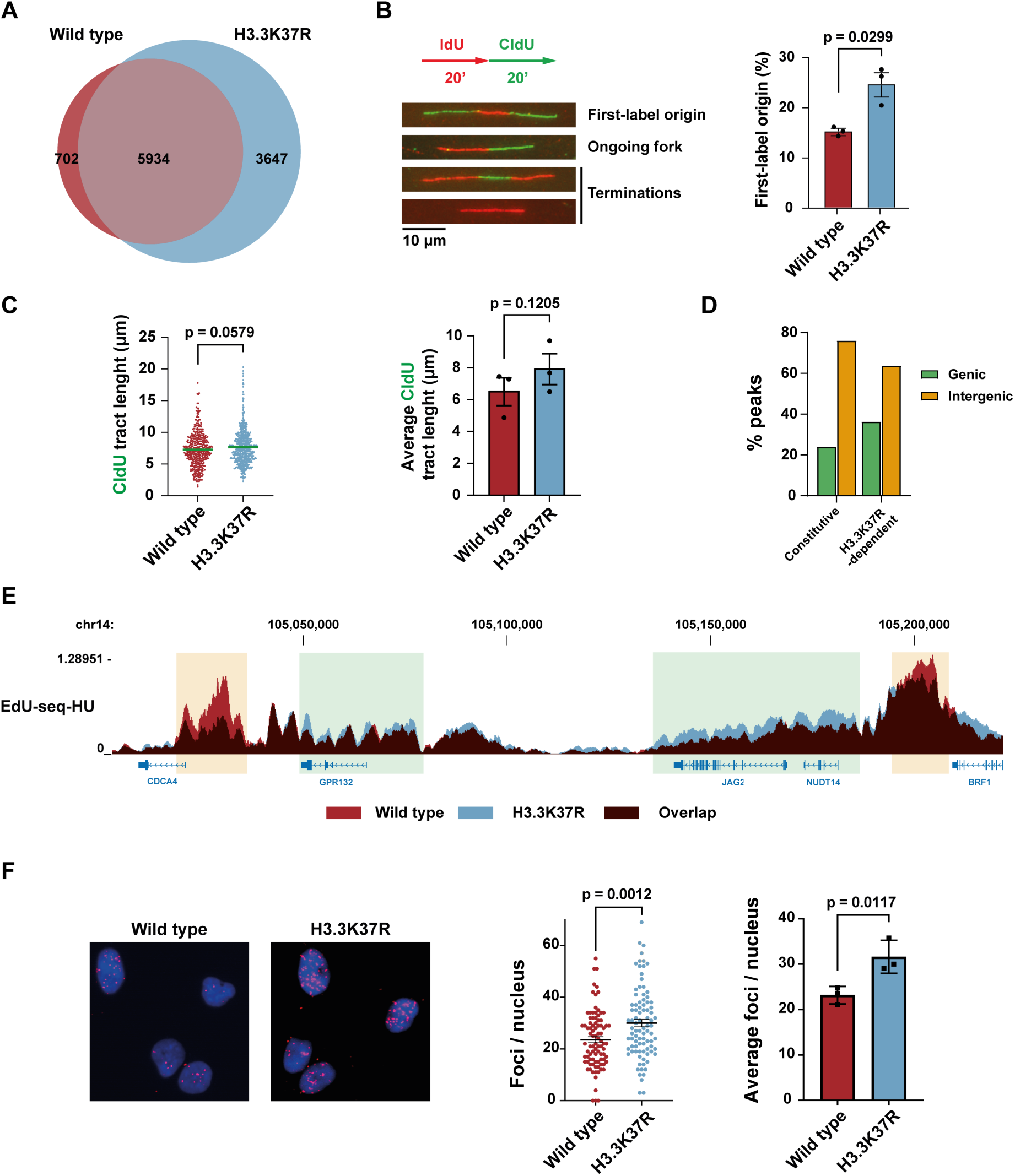
H3K37me1 restricts intragenic origin usage. **A.** Venn diagram showing unique and shared EdU-seq-HU peaks in wild-type and H3.3K37R mutant cells. **B.** Left: DNA replication patterns detected in stretched DNA fibers labeled with IdU and CldU. Right: Estimation of origin activity in wild-type and H3.3K37R mutant cells calculated as percentage of first-label origin structures. Data represent mean ± SEM of three independent experiments. **C.** Left: Representative distribution of fork speed values in wild-type and H3.3K37R mutant cells. (n = 466 and n = 615). Right: Average fork speed per independent experiment. Data represent mean ± SEM of three independent experiments. **D.** Distribution of EdU-seq–HU peaks in wild-type and H3.3K37R mutant cells with respect to gene annotation. **E.** Genome browser snapshot of EdU-seq-HU tracks in wild-type and H3.3K37R mutant cells. Green and yellow boxes indicate genic and intergenic regions, respectively. **F.** Left: Representative image of proximity-ligation assays (PLA) between RNA Pol II and PCNA in wild-type and H3.3K37R mutant cells. Middle: Representative distribution of RNA Pol II-PCNA PLA foci per nucleus in wild-type and H3.3K37R mutant cells (n = 90 and n = 90). Right: Average PLA foci per independent experiment. Data represent mean ± SEM of three independent experiments. Statistics by paired Student’s t test (B, C right and F right) or by unpaired Mann-Whitney test (C left and F middle).

To determine whether constitutive and H3.3K37R-induced EdUseq-HU peaks differed in their genomic distribution, we examined their localization relative to gene annotation. In agreement with previous reports,^15–18^ constitutive replication initiation sites preferentially localized to intergenic regions (Fig. 4D). However, a larger fraction of peaks gained in H3.3K37R cells mapped within protein-coding genes, pointing to a partial redistribution of initiation events toward intragenic regions upon loss of H3K37me1. Genome browser inspection of representative loci confirmed increased EdU incorporation within gene bodies in H3.3K37R mutant cells (Fig. 4E). If depletion of H3K37me1 increases intragenic origin firing, this would be expected to enhance the spatial proximity between transcription and replication machineries. To test this, we assessed RNA Pol II–PCNA proximity using proximity ligation assays (PLA). H3.3K37R cells exhibited significantly increased RNA Pol II–PCNA PLA signal (Fig. 4F and S4G), consistent with enhanced replication initiation within transcribed genes. Importantly, similar results were obtained using a second independent H3.3K37R mutant clone (Fig. S4H-I). Altogether, these results are consistent with a model in which H3K37me1 restricts intragenic origin usage.

### Loss of H3K37me1 leads to a redistribution of replication initiation activity

Increased EdU incorporation within gene bodies in H3.3K37R mutant cells was accompanied by reduced signal at intergenic initiation sites near TSSs (Fig. 4E), suggesting a redistribution of replication initiation activity. To systematically define this redistribution, we stratified protein-coding genes into two groups based on differential EdU incorporation between wild-type and H3.3K37R cells. Group I genes (n = 13090) displayed increased EdU-seq-HU signal across gene bodies while maintaining comparable TSS-proximal initiation levels (Fig. 5A), consistent with enhanced intragenic replication initiation. In contrast, Group II genes (n = 4823) exhibited an overall reduction in EdU-seq–HU signal (Fig. 5A), with a more pronounced decrease at TSS-proximal regions than within gene bodies. Importantly, though, both gene groups showed a significant increase in the gene-body/TSS ratio upon H3K37me1 loss (Fig. S5A), indicating a redistribution in the relative probability of replication initiation toward intragenic regions, independent of absolute initiation levels.

**Figure 5.**
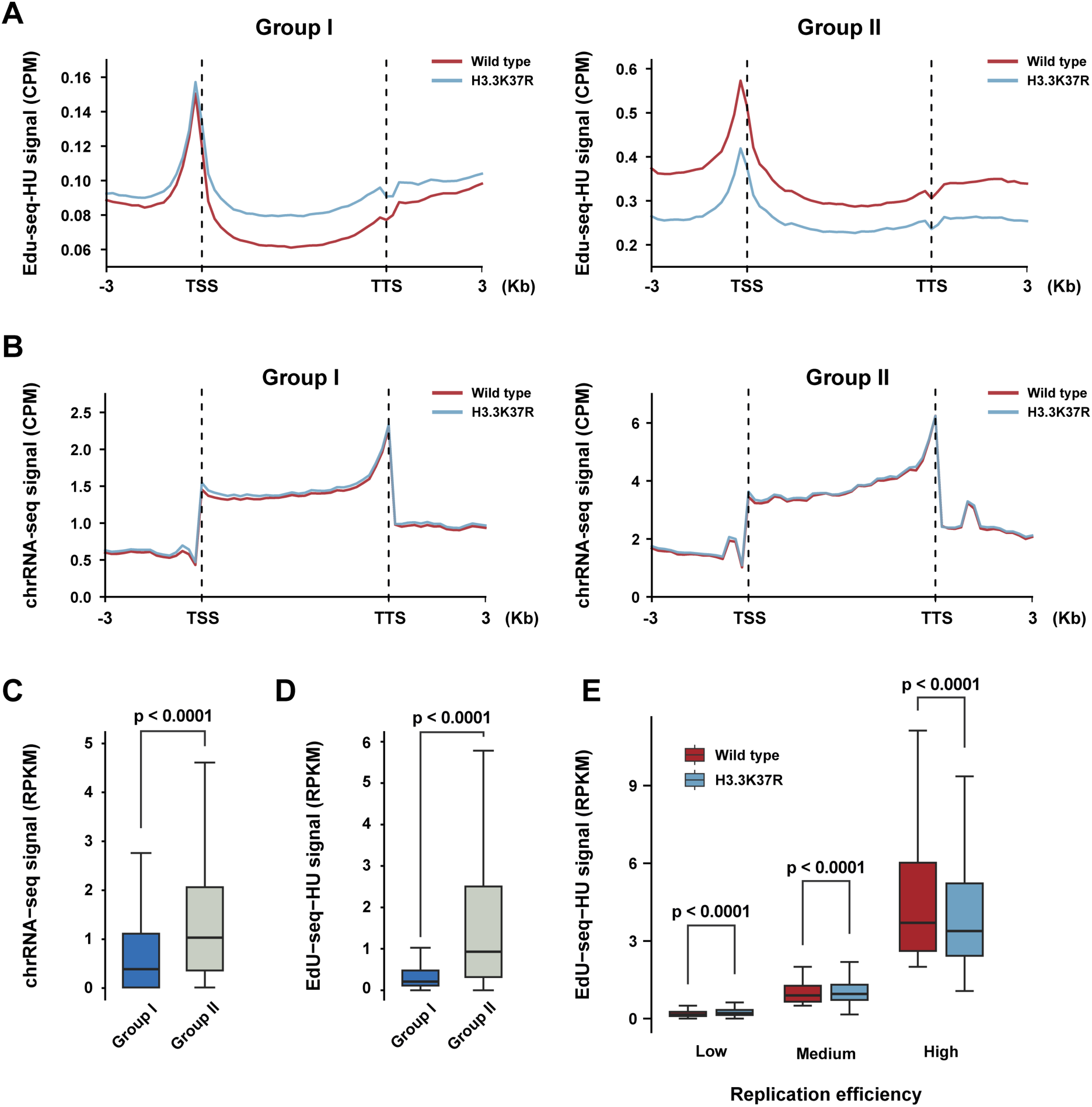
Loss of H3K37me1 leads to a redistribution of replication initiation activity. **A.** Metaplot of average EdU-seq–HU signal across Group I (left) and Group II (right) genes in wild-type and H3.3K37R mutant cells. **B.** Metaplot of average chrRNA-seq signal across Group I (left) and Group II (right) genes in wild-type and H3.3K37R mutant cells. **C.** Box plot of chrRNA-seq signal for Group I (left) and Group II (right) genes in wild-type cells. **D.** Box plot of EdU-seq–HU signal for Group I (left) and Group II (right) genes in wild-type cells. **E.** Box plot of EdU-seq–HU signal in wild-type and H3.3K37R mutant cells for genes plus 1 kb upstream stratified by initiation efficiency (low, mid, high). Statistics by paired Wilcoxon test (C, D, and E).

The redistribution of replication initiation activity observed upon H3K37me1 loss prompted us to investigate the basis of this locus-specific response. One possibility was that the distinct behaviors of Group I and Group II genes reflected differences in H3K37me1 depletion following H3.3K37R mutation. However, qChIP–seq analysis showed a comparable reduction of the mark across both gene categories in H3.3K37R cells (Fig. S5B). Thus, the different replication outcomes cannot be explained by locus-specific differences in H3K37me1 abundance.

Because transcription has been proposed to suppress intragenic origin licensing,^17,18^ we next asked whether the altered replication phenotype in H3.3K37R cells could arise indirectly from transcriptional perturbations. Metagene analysis of chrRNA-seq revealed highly similar transcriptional profiles between wild-type and mutant cells across both Group I and Group II genes (Fig. 5B), arguing against transcriptional deregulation as the primary driver of intragenic origin activation. Notably, however, the two gene categories differed intrinsically in their basal transcriptional output, with Group II genes exhibiting higher expression levels than Group I genes (Fig. 5C).

It is well-established that transcriptional activity positively correlates with origin efficiency and early replication timing, with highly transcribed loci displaying stronger initiation signals and preferential replication during early S phase.^15,24^ Consistent with this, Group II genes exhibited stronger EdU-seq–HU signal in wild-type cells (Fig. 5D), indicative of greater early-replication efficiency. Moreover, analysis of published Repli-seq datasets in RPE-1 cells ^24^ confirmed that these loci replicate preferentially during early S phase, whereas Group I genes exhibit comparatively later replication timing (Fig. S5C). To assess whether H3K37me1 loss differentially impacts loci according to their intrinsic initiation efficiency, we stratified genes based on EdU-seq–HU signal in wild-type cells. Genes with low or mid EdU-seq–HU signal exhibited increased initiation levels in H3.3K37R cells, whereas genes with the highest signal showed reduced initiation levels (Fig. 5E). Together, these findings suggest that loss of H3K37me1 leads to a shift in origin utilization from canonical early, efficient initiation sites toward intragenic sites that are normally inactive. This shift aligns with recent studies showing that recruitment of limiting replication initiation factors contributes to origin selection across the genome.^6,7^

### H3K37me1 limits MCM association with transcribed genes

Next, we sought to determine how H3K37me1 influences replication origin usage. It has been proposed that RNA Pol II–mediated transcription redistributes MCM2–7 complexes, thereby restricting early DNA replication initiation outside of active genes. ^17,18^ In this context, we hypothesized that transcription-associated H3K37me1 could restrict intragenic origin firing by limiting MCM2–7 chromatin association along transcribed regions. To test this hypothesis, we performed MCM2 qChIP–seq in wild-type and H3.3K37R cells. Consistent with previous reports,^18,19^ MCM2 was enriched at TSSs and relatively depleted across transcribed gene bodies in wild-type cells (Fig. S6A). Moreover, MCM2 levels across gene bodies inversely correlated with transcriptional output (Fig. S6A), in line with transcription-associated exclusion of MCM2–7 from actively transcribed regions. This distribution closely parallels the organization of early replication initiation detected by EdU-seq–HU (Fig. 5A). Importantly, loss of H3K37me1 significantly increased MCM2 occupancy across gene bodies (Fig. 6A-B). This intragenic accumulation was similarly observed across both Group I and Group II genes (Fig. 6C and S6B), suggesting that H3K37me1 broadly limits MCM2–7 association within transcribed regions. Together, these findings support a model in which H3K37me1 restricts MCM2–7 association within gene bodies, thereby limiting intragenic origin usage and preserving the canonical spatial organization of replication initiation.

**Figure 6.**
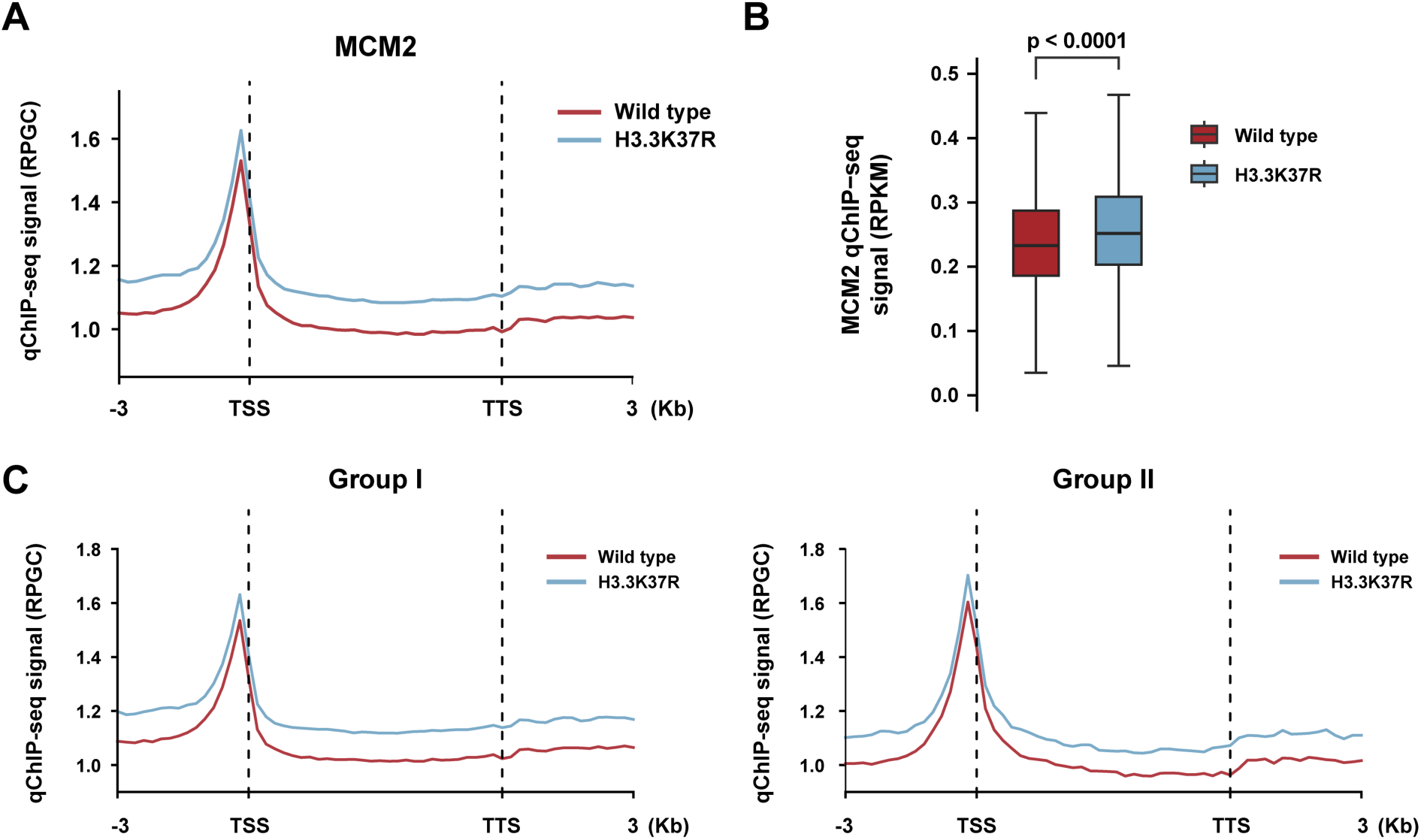
H3K37me1 limits MCM association with transcribed genes. **A.** Metaplot of average MCM2 qChIP-seq signal across protein-coding genes in wild-type and H3.3K37R mutant cells. **B.** Box plot of MCM2 qChIP-seq signal across protein-coding genes in wild-type and H3.3K37R mutant cells. **C.** Metaplot of average MCM2 qChIP-seq signal across Group I (left) and Group II (right) genes in wild-type and H3.3K37R mutant cells. Statistics by paired Wilcoxon test (B).

## Discussion

DNA replication initiation in metazoans occurs at thousands of potential sites distributed throughout the genome and follows a defined temporal program. Early replication initiation and active transcription both preferentially occur within open chromatin regions; however, the mechanisms that coordinate these two fundamental processes remain incompletely understood. Our work identifies H3K37me1, a transcription-associated histone modification, as an important determinant of the spatial organization of DNA replication initiation.

Although MCM2–7 loading is widespread, only a subset of licensed origins initiate replication, establishing a defined replication program across the genome. Early replication initiation sites are frequently located in non-transcribed regions adjacent to the TSS of actively transcribed genes, whereas replication initiation within gene bodies is generally suppressed.^15–18^ Prior studies have suggested that RNA Pol II–mediated transcription can reposition MCM2–7 complexes away from active genes,^17–19^ but the molecular basis for this redistribution has remained incompletely understood. Our results indicate that H3K37me1 contributes to this transcription-dependent reorganization, providing a chromatin-based mechanism linking transcriptional activity to replication origin usage.

Biochemical studies show that the MCM2 subunit directly interacts with a region of histone H3 spanning amino acid residues 26–67,^38^ raising the possibility that modifications within this segment influence MCM2–7 engagement with chromatin. Our observations support a model in which transcription-associated H3K37me1 helps to create a local chromatin environment that restricts MCM2–7 association within transcribed regions, thereby suppressing intragenic origin usage. Consistently, loss of H3K37me1 increases both MCM2–7 occupancy and replication initiation within gene bodies, indicating that H3K37me1 is required to maintain the spatial organization of DNA replication initiation in human cells.

We previously showed that H3K37me1 regulates DNA replication in *Saccharomyces cerevisiae*, where it is broadly distributed across the genome but relatively depleted at replication origins.^20^ However, unlike budding yeast, origin selection in higher eukaryotes is largely sequence-independent and occurs within broad initiation zones that are organized into a replication timing program. These initiation zones must operate in the context of diverse transcriptional states across different cell types.^39–42^ Consistent with this organization, our results indicate that H3K37me1 is selectively enriched across actively transcribed gene bodies in human cells. These observations suggest that H3K37me1 is integrated into a transcription-coupled chromatin mechanism that spatially organizes replication origin selection in metazoan genomes. By showing how origin usage can be regulated independently of DNA sequence, our work provides insight into the coordination of transcription with replication initiation in higher eukaryotes.

The close connection between transcription and DNA replication is supported by recent findings showing that the establishment of the replication timing program during early mammalian embryogenesis is preceded by, and partially dependent upon, transcriptional activation.^43^ In this setting, a histone modification-based mechanism that locally suppresses origin usage within transcribed regions may provide a flexible strategy to coordinate replication initiation with transcriptional activity without requiring fixed origin sequences. Such chromatin-driven control would help to minimize conflicts between the transcription and replication machineries while preserving the integrity of the DNA replication program across differentiation.

Our work supports the emerging view that the DNA replication landscape is shaped not only by pathways that activate origins but also by repressive mechanisms that locally restrict replication initiation.^44–46^ In this context, histone PTMs may function not only as markers of individual DNA-templated processes but also as regulatory signals that help to integrate their activities,^47^ thereby facilitating communication between transcription and DNA replication to coordinate genome function and maintenance over time.

## Methods

### Cell culture

hTERT RPE-1 cells (American Type Culture Collection, ref. CRL-4000) were cultured in Dulbecco’s Modified Eagle’s Medium/Nutrient Mixture F-12 Ham (Sigma-Aldrich, ref. D8437-500ML) supplemented with 10% fetal bovine serum (Sigma-Aldrich, ref. F7524) and 100 U/mL penicillin / 100 µg/mL streptomycin (Corning, ref. 702633). Cells were grown in a Forma™ Steri-Cycle™ CO_2_ Incubator (ThermoFisher Scientific, ref. 371) at 37°C and 5% CO_2_. Mycoplasm test was conducted regularly.

### CRISPR Editing

For generation of the H3.3B K37R clones, 1.5×10^6^ hTERT-RPE1-Cas9 cells were transfected with 30 pmol of ssODN template (99nt, Alt-R™ HDR Donor Oligo, Integrated DNA technologies, sequence: GGCCACGAAAGCCGCCAGGAAAAGCGCTCCCTCTACCGGCGGTGTGAAGCGGCCTCATCGCTACAGGTAGGTCGGGCGGGGGAACAATGGCCCGGCGGT) and 80 pmol H3.3B sgRNA (Alt-R^®^ CRISPR-Cas9 crRNA and tracRNA, sequence: AAGCGCTCCCTCTACCGGCG) using Lipofectamine RNAiMAX Transfection Reagent (Invitrogen), following manufacturer’s instructions for reverse transfection. For H3.3A KO mutant clones, cells were transfected with the H3.3A sgRNA only (sequence: TTTTAATACCTGTAACGATG). Cells were incubated 1 h prior to lipofection and 8 h after lipofection with 1 µM of the DNA-PK inhibitor NU7441 and 10 µM of the polymerase θ inhibitor ART558 to increase gene editing efficiency. Single cell sorting was carried out 72 h after transfection into conditioned media. Colonies were expanded for genotyping and freezing. The H3.3B K37R lines were generated from previously generated H3.3 KO lines.

For screening H3.3 KO clones, DNA was extracted using direct lysis buffer (10 mM Tris pH 8.0, 2.5 mM EDTA, 0.2 M NaCl, 0.15% SDS, 0.3% Tween-20). The lysate was heated to 97°C for 60 s on a thermocycler to ensure lysis, then amplified using the MyTaq™ DNA Polymerase (Bioline) following manufacturer’s instructions and sequenced by Sanger sequencing. Screening for H3.3B K37R clones was carried out based on the method described in ^48^ with the following modifications: for DNA extraction from 96 wells plates, cells were collected using 25 µL trypsin, 5 µL of which was transferred to fresh 96-well PCR plate containing 15 µL direct lysis buffer. The lysate was heated to 97°C for 60 s on a thermocycler to ensure lysis, then diluted 1:3 in ddH_2_O. Diluted lysate was used in a 10 µL qPCR reaction using iTAQ 2x supermix (BioRad). Primers were designed to only amplify from successful edits; H3.3B mutant sequences fwd: GTGGGGCCAGGATCTGTTC and rev: CTGTAGCGATGAGGCCGC. Positive clones were confirmed by Sanger sequencing with the following primers: H3.3A fwd: GGTGATCGTGGCAGGAAAAG, H3.3A rev: CCAGGTAAGATTATGGCTTCAAAGT, H3.3B fwd: CCTTATCTTCGGGGCGTCTT and H3.3B rev: TCCAGGCCTTTGTCTTACCT.

### Histone extraction and digestion

Histone proteins were extracted from as described by ^49,50^. Briefly, histones were acid-extracted with chilled 0.2 M sulfuric acid (5:1, sulfuric acid : pellet) and incubated with constant rotation for 4 h at 4°C, followed by precipitation with 33% trichloroacetic acid (TCA) for 1 h at 4°C. Then, the supernatant was removed and the tubes were rinsed with ice-cold acetone containing 0.1% HCl, centrifuged and rinsed again using 100% ice-cold acetone. After the final centrifugation, the supernatant was discarded and the pellet was dried at room temperature. The pellet was dissolved in 50 mM ammonium bicarbonate, pH 8.0, and histones were subjected to derivatization using 5 µL of propionic anhydride and 14 µL of ammonium hydroxide (all Sigma Aldrich) to balance the pH at 8.0. The mixture was incubated for 15 min and the procedure was repeated. Histones were then digested with 1 µg of sequencing grade trypsin (Promega) diluted in 50mM ammonium bicarbonate (1:20, enzyme:sample) overnight at room temperature. Derivatization reaction was repeated to derivatize peptide N-termini. The samples were dried in a vacuum centrifuge.

### LC-MS/MS Acquisition and Analysis

Prior to mass spectrometry analysis, samples were desalted using a 96-well plate filter (Orochem) packed with 1 mg of Oasis HLB C-18 resin (Waters). Briefly, the samples were resuspended in 100 µL of 0.1% TFA and loaded onto the HLB resin, which was previously equilibrated using 100 µL of the same buffer. After washing with 100 µL of 0.1% TFA, the samples were eluted with a buffer containing 70 µL of 60% acetonitrile and 0.1% TFA and then dried in a vacuum centrifuge.

Samples were resuspended in 10 µL of 0.1% TFA and loaded onto a Dionex RSLC Ultimate 300 (Thermo Scientific), coupled online with an Orbitrap Fusion Lumos (Thermo Scientific). Chromatographic separation was performed with a two-column system, consisting of a C-18 trap cartridge (300 µm ID, 5 mm length) and a picofrit analytical column (75 µm ID, 25 cm length) packed in-house with reversed-phase Repro-Sil Pur C18-AQ 3 µm resin. Peptides were separated using a 30 min gradient from 1-30% buffer B (buffer A: 0.1% formic acid, buffer B: 80% acetonitrile + 0.1% formic acid) at a flow rate of 300 nL/min. The mass spectrometer was set to acquire spectra in a data-independent acquisition (DIA) mode. Briefly, the full MS scan was set to 300-1100 m/z in the orbitrap with a resolution of 120,000 (at 200 m/z) and an AGC target of 5×10^5^. MS/MS was performed in the orbitrap with sequential isolation windows of 50 m/z with an AGC target of 2×10^5^ and an HCD collision energy of 30.

Histone peptides raw files were imported into EpiProfile 2.0 software.^51^ From the extracted ion chromatogram, the area under the curve was obtained and used to estimate the abundance of each peptide. In order to achieve the relative abundance of post-translational modifications (PTMs), the sum of all different modified forms of a histone peptide was considered as 100% and the area of the particular peptide was divided by the total area for that histone peptide in all of its modified forms. The relative ratio of two isobaric forms was estimated by averaging the ratio for each fragment ion with different mass between the two species. The resulting peptide lists generated by EpiProfile were exported to Microsoft Excel and further processed for a detailed analysis. Log_2_ fold change and p-value for H3.3K37R-wild type comparison were determined using custom scripts under R 4.2.3 for volcano plots.

### DNA fiber analysis

Cells were pulse-labeled with 20 µM IdU (20 min) followed by 200 µM CldU (20 min), prior to cell harvesting by scrapping in phosphate-buffered saline (PBS) + 0.1% bovine serum albumin (BSA). Cells were then pelleted and resuspended in PBS + 0.1% BSA at a final concentration of 1-2×10^3^ cells/µL. 2.5 µL of cell suspension were spotted on a positively charged slide and lysed with 7.5 µL of spreading buffer (200 mM Tris-HCl pH 7.5, 50 mM EDTA, 0.5% SDS). After 8 min, slides were tilted at 45 degrees to allow the DNA to spread. Slides were then air-dried, fixed with ice-cold methanol/acetic acid (3:1) for 5 mins, air-dried and stored at 4°C. Slides were rehydrated with PBS, denatured with 2.5 M HCl for 1h, washed with PBS twice, and blocked with blocking buffer (3% BSA, 0.1% Triton X-100 in PBS) for 40 min. Next, slides were incubated with primary antibody mix of mouse anti-BrdU which recognizes IdU (1:250, Becton Dickinson, ref. 347580) and rat anti-BrdU which recognizes CldU (1:250, Abcam, ref. 6326) diluted in blocking buffer for 2.5 h at RT in a dark humid chamber. Slides were washed 3 times with PBS for 5 min each and incubated with secondary antibodies anti-mouse Alexa fluor 594 and anti-rat Alexa fluor 488 (1:250, Invitrogen, ref. A11032 and ref. A11006, respectively) in blocking buffer for 1h at RT in a dark humid chamber. After washing 3 times with PBS and air-drying, slides were mounted with Prolong gold antifade reagent (Invitrogen, ref. P36930) and stored at 4°C until imaging. Images were acquired using a AF6000 Leica fluorescence microscope or a DMi8 Leica Thunder fluorescence microscope.

For the CldU tract length analysis, at least 350 fibers were measured per condition in each independent experiment using the segmented line tool on ImageJ FIJI software (https://fiji.sc). For origin firing analysis, origins labeled during the first pulse (green-red-green structures) were quantified as percentage of all structures containing red (>450 total structures scored per condition in each independent experiment). Graphics and statistics were generated within GraphPad 10.6.1 software.

### In situ proximity ligation assays (PLA)

5×10^4^ cells were seeded in 12-well plates, pre-permeabilised in CSK buffer (10 mM PIPES pH 6.8, 0.1 M NaCl, 0.3 M sucrose, 0.25% Triton X-100) for 3 min, fixed in paraformaldehyde-PBS 4% for 10 min at room temperature and permeabilised in Triton X-100-PBS 0.5% for 5 min. Then, PLA protocol from sigma-Aldrich was followed, using anti-mouse MINUS (DUO92004) and anti-rabbit PLUS (DUO92002) probes, and Duolink Detection Reagents (DUO92008). Cells were counterstained with DAPI and mounted in VECTASHIELD mounting medium from Vector Laboratories. Images were captured within a DM6000B Leica fluorescence microscope.

PLA foci were counted using custom scripts in ImageJ FIJI software. Graphics and statistics were generated within GraphPad 10.6.1 software.

### Flow cytometry

3×10^5^ cells were seeded into 6 cm plates. Samples were fixed with 90% methanol and incubated overnight at -20°C. Then, cells were washed with ice-cold PBS + 1% BSA and permeabilised with PBS + 0.2% Triton X-100. After two more washes with PBS + 1% BSA, EdU Click-It reaction was carried out for 30 min at room temperature using the following mix: 100 mM Tris-HCl pH 8.5, 1 mM CuSO_4_, 1 µM Alexa Fluor Azide (Invitrogen, ref. A10277) and 100 mM ascorbid acid. Finally, cells were resuspended in PBS + 10 µg/mL propidium iodide (Sigma, ref. P4170) + 20 µg/mL RNAse A (Roche, ref. 10109169001). Cytometry was carried out within a BD FACSCalibur and analysed using FlowJo 10.10.0.

### EdU-seq-HU

Experiments were carried out as described in ^35^ with the following modifications: to stall cells at the G1/S checkpoint, prior to EdU/HU incubation, cells were seeded at 50% confluency then incubated in 0.5 µM palbociclib (Selleck Chemicals, ref. S1116) for 24 h. After 24 h cells were washed in warmed PBS and cell media, incubated for 5 h in fresh media containing 25 µM EdU (Sigma, ref. 900584) and 2 mM HU (Sigma-Aldrich, ref. H8627), then collected using Trypsin. 5×10^6^ cells were collected per replicate. For DNA extraction, cells were incubated in lysis buffer overnight at 50°C and 700 RPM. For library preparation, following isolation of EdU-labelled DNA by Dynabeads MyOne Streptavidin C1 beads, on-bead tagmentation was carried out. Briefly, beads were washed in 10mM Tris-HCl pH 8 and resuspended in tagmentation buffer (10µL 5X Tag buffer (50mM Tris-HCl pH 8, 50% DMF, 25mM MgCl_2_), 2 µL of tagmentase, 38µL water), then incubated for 10mins at 37°C. The reaction was inhibited on ice for 2 mins. Beads were then washed in ice cold wash buffer (20 mM Tris-HCl pH 8, 250 mM LiCl, 1 mM EDTA, 1% NP-40, 1% Sodium deoxycholate) and transferred to Lo-Bind 1.5ml tubes (Eppendorf). Finally, beads were washed in 10mM tris-HCL pH8 and transferred to PCR tubes. Sequencing libraries were prepared using the NEBnext 2x Master Mix (New England Biolabs, ref. M0541S) via on-bead PCR. Final libraries were size selected using Ampure XP beads (Beckman Coulter).

### Chromatin immunoprecipitation sequencing

A total of 1×10^6^ cells (or 3×10^6^ cells for MCM2 ChIP-seq) were collected with trypsin (Corning, ref. 15333651) and crosslinked using 1% formaldehyde (Sigma-Aldrich, ref. 252549) in cell medium for 13 min at 37°C. Sonication was conducted within a Covaris^®^ E220*evolution* Focused-ultrasonicator. Nuclei fraction was isolated according to manufacturer’s instructions. Protease activity was inhibited by adding cOmplete (Roche, ref. 11873580001) and PMSF (Millipore, ref. 52332) to lysis, wash and shearing buffers. Sonication time was previously optimized between 15 and 30 min while the rest of parameters remained as recommended. Sheared chromatin was then clarified by centrifuging for 10 min at 13000 RPM and 4°C.

Chromatin was diluted in IP buffer (20 mM TrisHCl pH 8, 150 mM NaCl, 2 mM EDTA, 1% Triton X-100, 0.1% SDS) and incubated overnight at 4°C in a tube rotator with the corresponding antibody: H3.3 (Merck, ref. 09-838), H3K36me1 (Cell Signalling, ref. 14111S), H3K36me3 (Thermo Fisher ref. MA5-24687), H3K37me1 (Abcam, ref. AB215728) and MCM2 (Cell Signalling, ref. 3619S). For qChIP-seq samples, sheared chromatin from NMuMG cells was added at a 1:20 ratio. Afterwards, 25 µL Dynabeads Protein A (Thermo Fisher, ref. 10001D) were pre-washed and added to chromatin for a 2 h incubation in rotation at 4°C. Chromatin was washed twice with IP buffer, twice with Wash buffer (20 mM TrisHCl pH 8, 500 mM NaCl, 2 mM EDTA, 1% Triton X-100, 0.1% SDS) and twice with LiCl buffer (20 mM TrisHCl pH 8, 250 mM LiCl, 1 mM EDTA, 1% IGEPAL/NP-40, 1% Na deoxycholate). Then tagmentation was performed as described for EdU-seq-HU section, and samples were washed again with LiCl buffer and twice with 1x TE pH 8 (10 mM Tris-HCl, 1 mM EDTA). DNA was eluted from beads through an incubation in Elution buffer (0.1 NaHCO_3_, 1% SDS) for 30 min at 65°C. In order to reverse the crosslinking, DNA was incubated overnight at 65°C after adding 0.5 M NaCl and 0.5 mg/mL Proteinase K (Invitrogen, ref. 11588916). RNA was removed by adding 0.1 mg/mL RNAse A (Thermo Scientific, ref. EN0531) and incubating for 1 h at 37°C. Following purification of DNA was carried out using the ChIP DNA Clean & Concentrator kit (ZYMO, ref. D5205). Inputs were tagmentated as before and then purified using Ampure Beads. Sequencing libraries were prepared using the NEBnext 2x Master Mix. Final library size selection was carried out as in EdU-seq-HU section.

### Chromatin associated RNA sequencing

A total of 4×10^6^ cells were collected and isolation of chromatin-associated RNA was performed as described in ^52^. Final chromatin fraction RNA was extracted using TRIzol (Thermo Scientific, ref. 15596026) and the Direct-zol RNA MiniPrep Kit (Zymo Research, ref. R2050). Libraries were prepared at the CABIMER Genomics Facility (CABIMER, Sevilla) with the Illumina Stranded Total Prep with Ribo Zero Plus under manufacturer’s protocol recommendations.

### High-throughput sequencing and data processing

Libraries were quantified using the Qubit HS kit (Invitrogen) and analysed for size profiles with Bioanalyzer. Sequencing was performed at the CABIMER Genomics Facility (CABIMER, Sevilla) in an Illumina NextSeq 500 High-Output with 1x 75 bp single-end configuration or in an Illumina Novaseq 6000 system with 2x 50 bp paired-end configuration. Then, samples were de-multiplexed and quality filtered, and adaptors were removed.

Raw reads from newly generated ChIP-seq and EdU-seq-HU samples and downloaded ChIP-seq, ATAC-seq and Repli-seq samples were aligned using the Burrows-Wheeler Aligner^53^ (BWA) v0.7.17-r1188 (-n 3 -k 2 -R 300) to the Genome Reference Consortium Human build 38 (GRCh38/hg38). Multimappers were removed with samtools ^54^ v1.13 view (-q 30 -F 780) as well as duplicates with samtools markdup (default parameters). qChIP-seq samples were also mapped to the Genome Reference Consortium Mouse build 39 (GRCm39/mm39) and processed with the same parameters. Similarly, raw reads from chrRNA-seq and downloaded EU-seq (nascent RNA) samples were aligned using Spliced Transcripts Alignment to a Reference ^55^ (STAR) v2.7.11b (default parameters) and filtered with samtools but no duplicates were discarded.

The next processing steps were performed under deeptools v3.5.6.^56^ Genome coverage was calculated using bamCoverage at 50 bp bin size resolution. ChIP-seq simples were normalized in reads per genome coverage (RPGC) and scaled with the occupancy ratio described in ^57^ when spike-in chromatin was added. After validating the reproducibility, independent replicates (n = 2) were merged for visualization and further analyses.

Quantification of reads was assessed using multiBamSummary from deeptools v3.5.6 ^56^. Annotated genes were previously downloaded from Biomart (Ensembl, release 112), filtering for protein coding genes on chromosomes 1-22. Outputted raw counts were then normalized in reads per kilobase and millions of reads (RPKM) to avoid gene length bias on signal estimation. Solely, quantification of EdU-seq-HU at 50 Kb bin resolution was normalized in CPM. Additionally, qChIP-seq simples were scaled by the same OR as described before for genome coverage tracks. At this point, mean between replicates was computed before creating scatterplots and boxplots with custom R scripts. Difference between signal medians were considered significant when p-value < 0.05 (paired Wilcoxon test).

### Peak calling

H3K37me1 and EdU-labelled enriched regions were identified using MACS3^58^ v3.0.2 callpeak with -q 0.01 --nomodel --nolambda parameters. In addition, minimum length of 5 Kb and a maximum distance between peaks of 10 Kb (--min-length 5000 --max-gap 10000) were established accounting for broad H3K37me1-positive regions rather than narrow peaks. For EdU peaks, these parameters were reduced to --min-length 200 --max-gap 1000. Finally, only H3K37me1 and EdU peaks displaying > 3 and > 7 fold enrichment respectively were considered for further analysis.

### Differential expression and gene ontology analysis

chrRNA read counts in protein coding genes were computed using featureCounts^59^ v2.0.3 regarding on reverse strand specificity (-s 2), paired-end layout (--countReadPairs) and overlap between features (-O). Differential expressed genes were determined using DESeq2^60^ v1.40.0 (|log_2_ fold change| > 2 and q-value < 0.05) under R V4.3.0. Gene ontology analysis results and graphics were generated within R package clusterProfiler^61^ v4.10.0. Only biological processes with an adjusted p-value < 0.05 were considered.

## Acknowledgments

We thank the Genomic and Microscopy Core facilities at CABIMER for their technical help, and Maria Elena Torres-Padilla and Belén Gómez-González for critically reading the manuscript. We also thank Dr. Daniel Durocher for generously providing the RPE-1 Cas9 cell line. The G.M.-Z. laboratory was supported by grants PID2021-127432NA-I00 and PID2023-151942NB-I00 funded by MICIU/AEI/10.13039/501100011033 and by ERDF/EU, by grant CNS2022-135600 funded by MICIU/AEI/10.13039/501100011033 and by European Union NextGenerationEU/PRTR, and by the EMERGIA 2021 program from the Junta de Andalucía (EMC21_00222). The N.G.-R. laboratory was supported by grant PID2023-150072N, funded by MICIU/AEI/10.13039/501100011033 and by ERDF/EU; by grant CNS2024-154523 funded by MICIU/AEI/10.13039/501100011033; and by the EMERGIA 2021 program from the Junta de Andalucía (EMC21_00057). The Sidoli lab gratefully acknowledges for funding the Hevolution Foundation (AFAR), the Einstein-Mount Sinai Diabetes center, the ERCM-CFAR center for AIDS research, and the NIH Office of the Director (S10OD030286). R.A. was supported by the University of Seville (PIF-2023). D.P.-A. was supported the Ministry of Science, Innovation and Universities (FPU23/03088). G.M.-Z. was supported by a fellowship from “la Caixa” Foundation (ID 100010434) and from the European Union’s Horizon 2020 research and innovation programme under Marie Skłodowska-Curie grant agreement No. 847648. The fellowship code is LCF/BQ/PR21/11840007.

## Author contributions

Conceptualization: G.M.-Z.; Data curation: R.A., L.L.-H., S.S. and L.P.; Formal analysis: R.A., L.L.-H., S.S. and L.P.; Funding acquisition: S.S., N.G.-R. and G.M.-Z.; Investigation: R.A., P.T.-K., I.D.-G. and D.P.-A.; Project administration: G.M.-Z.; Software: R.A., L.L.-H. and L.P.; Supervision: N.G.-R. and G.M.-Z.; Visualization: R.A. and I.D.-G.; Writing – original draft: G.M.-Z. Writing – review & editing: S.S., N.G.-R., L.P. and G.M.-Z.

---

**Figure S1.**
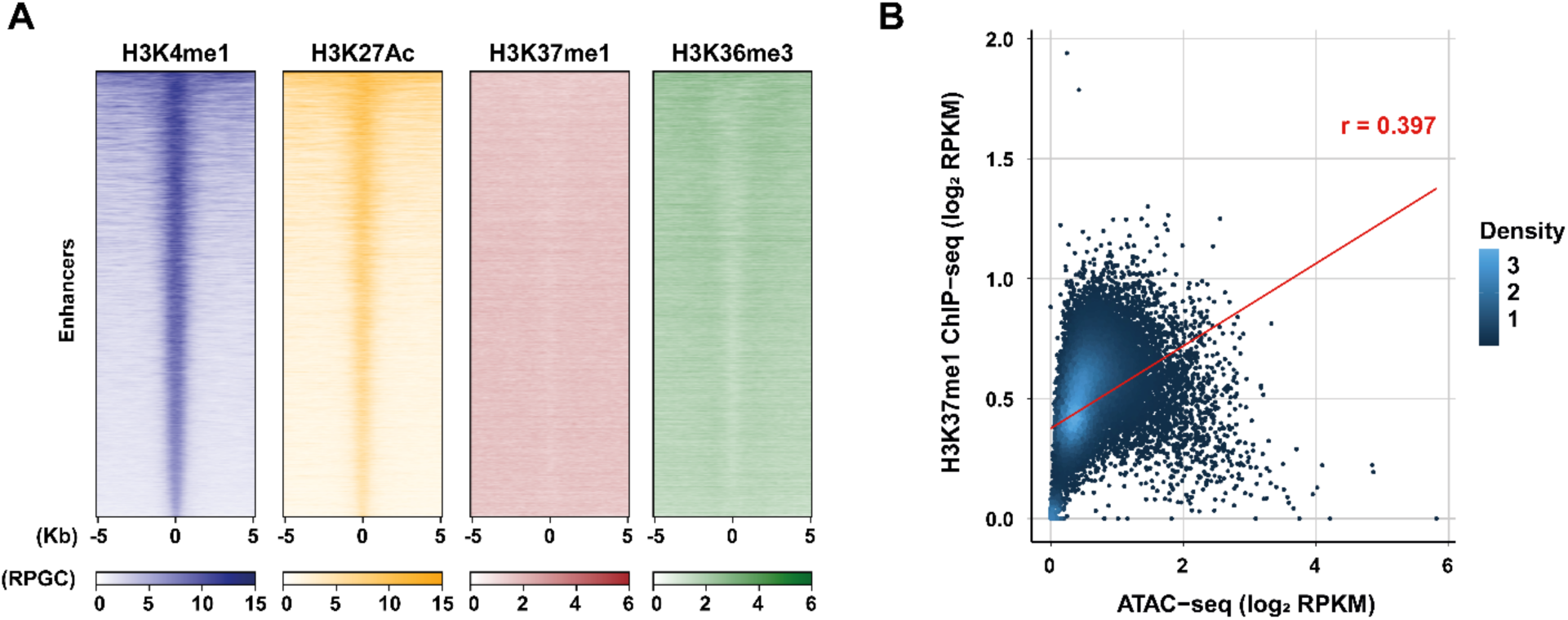
Supplementary to H3K37me1 marks actively transcribed gene bodies. **A.** Heatmaps of H3K4me1, H3K27ac, H3K37me1 and H3K36me3 ChIP-seq signal across enhancers. Rows are sorted according to total H3K4me1 ChIP-seq intensity. **B.** Scatterplot of ATAC-seq (x-axis) versus H3K37me1 ChIP-seq (y-axis) signal at annotated genes.

**Figure S2.**
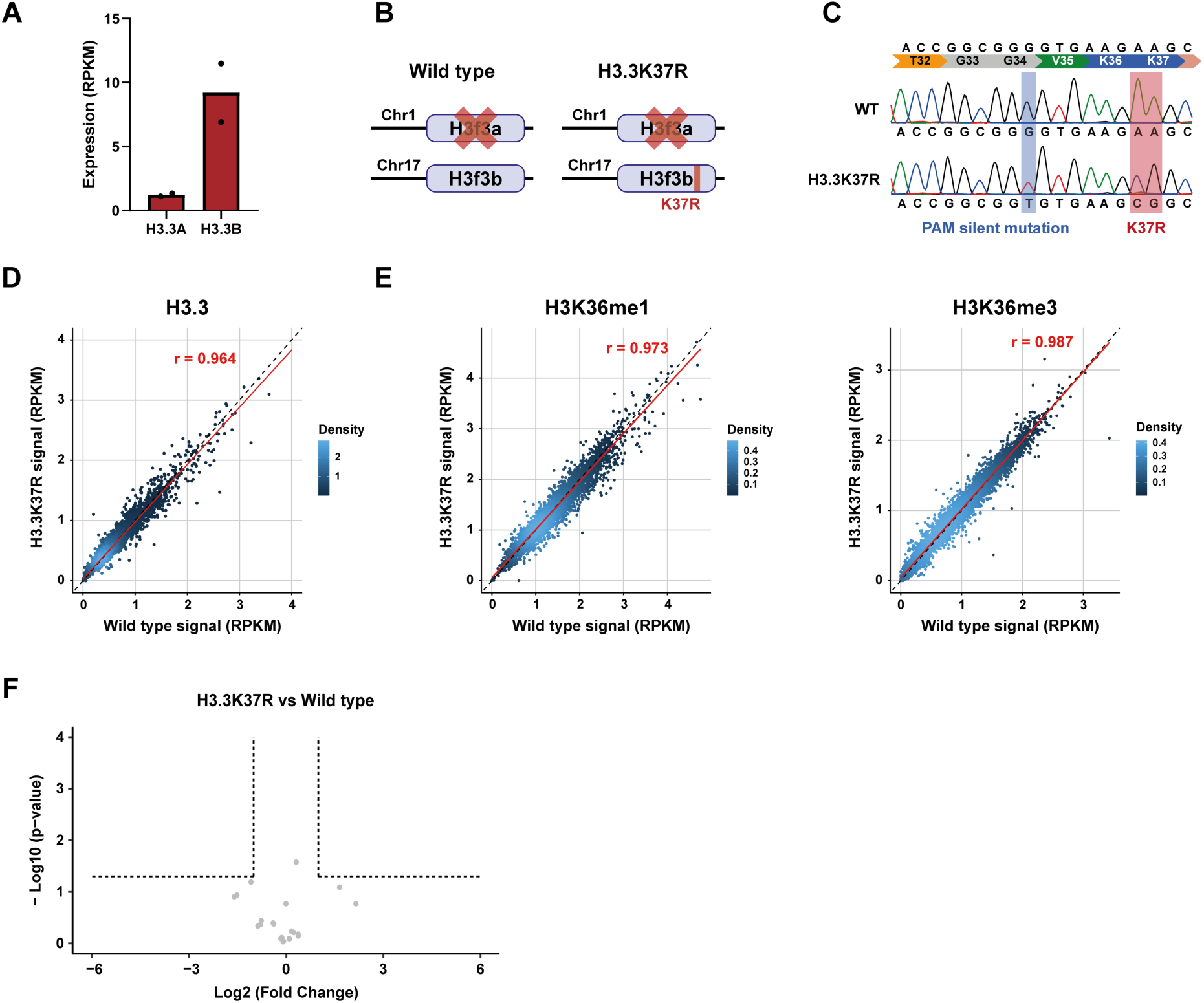
Supplementary to H3.3K37R mutation selectively reduces H3K37me1 levels. **A.** Expression levels (RPKM) of H3.3A and H3.3B mRNAs in RPE-1 cells. **B.** Schematic of the wild-type (H3F3A–/–; H3F3BWT/WT) and H3.3K37R mutant (H3F3A–/–; H3F3BK37R/K37R) genotypes. **C.** Chromatograms from Sanger sequencing in wild-type and H3.3K37R mutant cells. **D.** Scatterplot comparing H3.3 qChIP-seq signal across euchromatic genes in wild-type (x-axis) and H3.3K37R (y-axis) mutant cells. **E.** Scatterplots comparing H3K36me1 (left) or H3K36me3 (right) qChIP-seq signals across euchromatic genes in wild-type (x-axis) and H3.3K37R (y-axis) mutant cells. **F.** Volcano plots displaying global changes in histone H4 post-translational modifications detected by mass spectrometry analysis in H3.3K37R versus wild-type cells. Two-sided Student’s t-test; n = 3 independent replicates; |log_2_ FC| > 1; p-value < 0.05.

**Figure S3.**
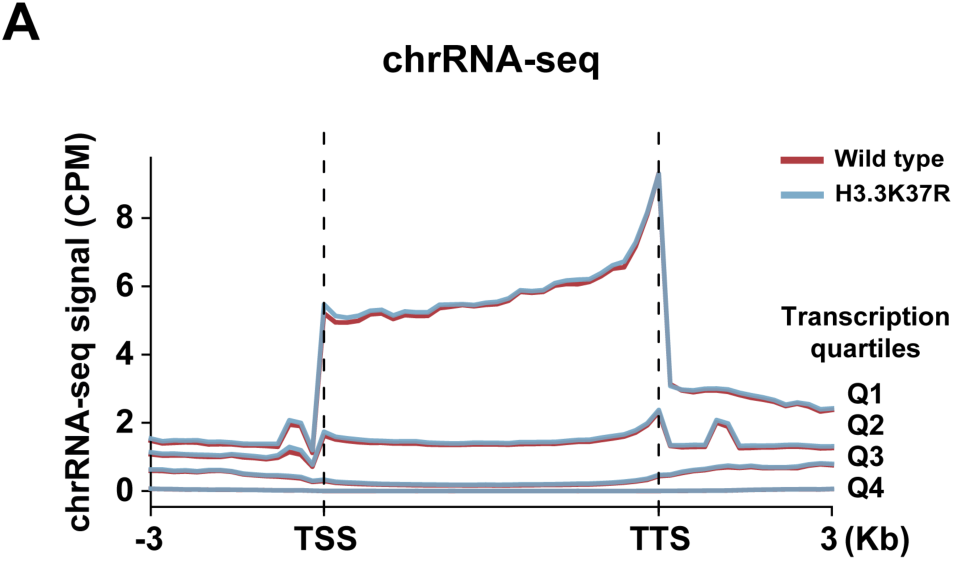
Supplementary to loss of H3K37me1 does not globally alter transcription. **A.** Metaplot of average chrRNA-seq signal in wild-type and H3.3K37R mutant cells along genes grouped into quartiles (Q1–Q4) by transcription level.

**Figure S4.**
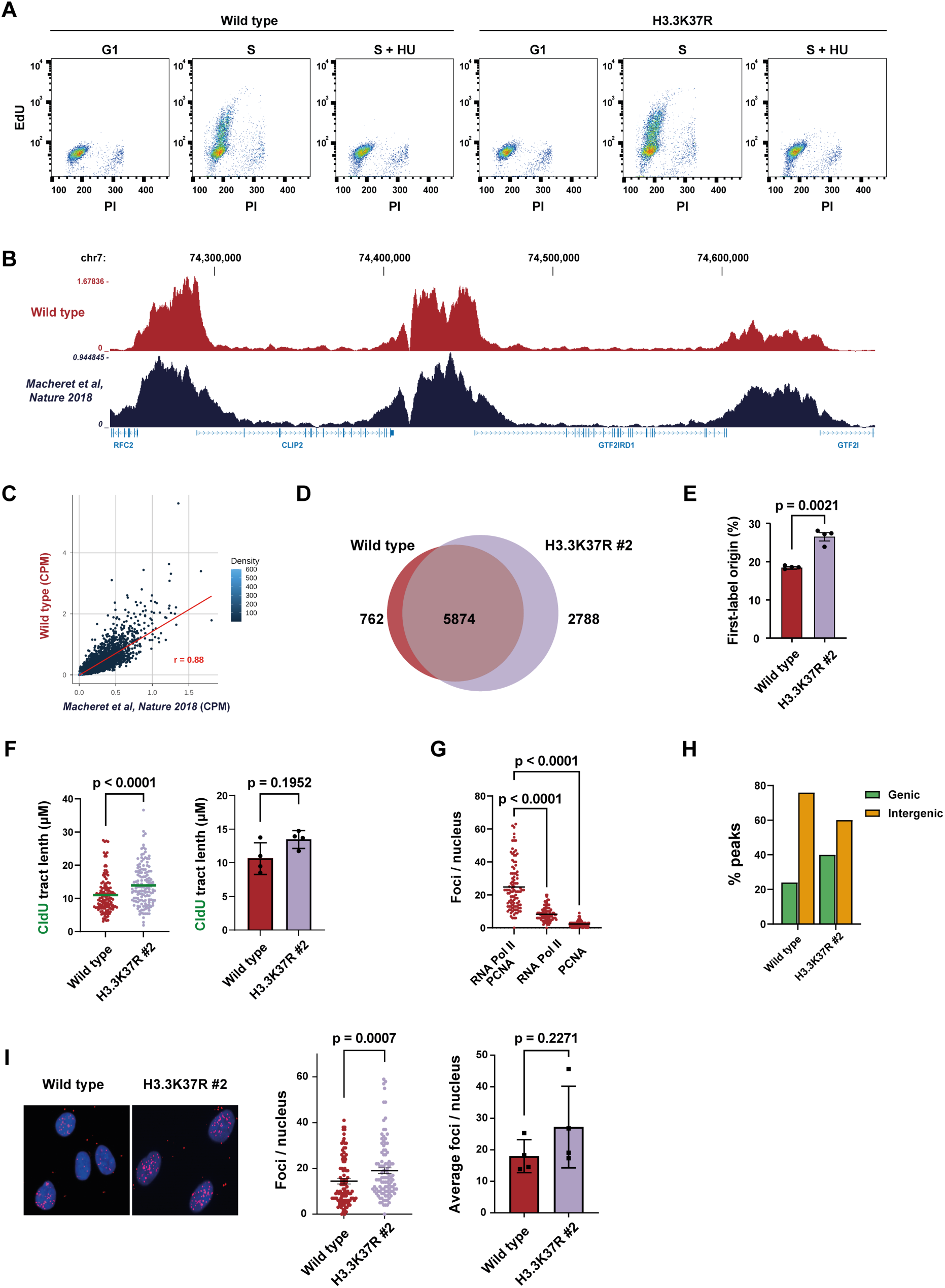
Supplementary to H3K37me1 restricts intragenic origin usage. **A.** EdU-FACS analysis of G1-arrested cells at 0 h (G1) and 5 h after release into S phase in the absence (S) or presence (S + H) of 2 mM HU. **B.** Genome browser snapshot of EdU-seq-HU tracks in wild-type cells (this study) and in RPE-1 cells from *Macheret et al* ^17^. **C.** Scatterplot comparing EdU-seq-HU signal (50kb bins) in RPE-1 cells from *Macheret et al* ^17^ (x-axis) and wild-type cells from this study (y-axis). **D.** Venn diagram showing unique and shared EdU-seq-HU peaks in wild-type and H3.3K37R #2 mutant cells. **E.** Estimation of origin activity in wild-type and H3.3K37R #2 mutant cells calculated as percentage of first-label origin structures. Data represent mean ± SEM of four independent experiments. **F.** Left: Representative distribution of fork speed values in wild-type and H3.3K37R mutant cells. (n = 136 and n = 145). Right: Average fork speed per independent experiment. Data represent mean ± SEM of four independent experiments. **G.** Control experiment for the PLA assay showing distributions of PLA foci per nucleus obtained with RNA Pol II only, PCNA only, or both antibodies together in wild-type cells. **H.** Distribution of EdU-seq–HU peaks in wild-type and H3.3K37R #2 mutant cells with respect to gene annotation. **I.** Left: Representative image of proximity-ligation assays (PLA) between RNA Pol II and PCNA in wild-type and H3.3K37R mutant cells. Middle: Representative distribution of RNA Pol II-PCNA PLA foci per nucleus in wild-type and H3.3K37R mutant cells (n = 104 and n = 104). Right: Average PLA foci per independent experiment. Data represent mean ± SEM of four independent experiments (paired t-test). Statistics by paired Student’s t test (E, F right and I right) or by unpaired Mann-Whitney test (F left, G and I middle).

**Figure S5.**
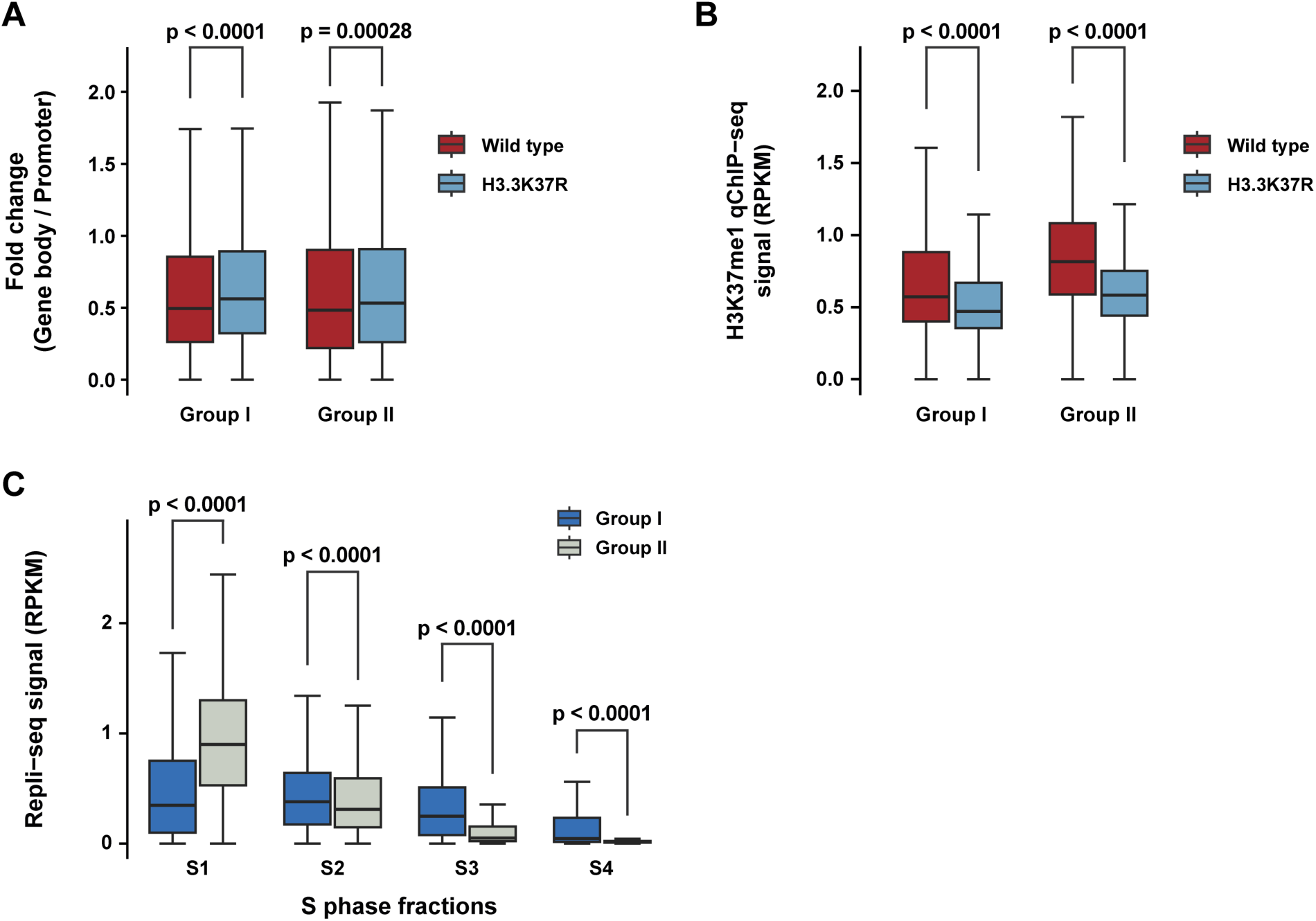
Supplementary to loss of H3K37me1 leads to a redistribution of replication initiation activity. **A.** Box plot of gene-body/TSS EdU-seq–HU ratios for Group I and II genes in wild-type and H3.3K37R mutant cells. **B.** Box plot of H3K37me1 qChIP-seq signal for Group I and Group II genes in wild-type and H3.3K37R mutant cells. **C.** Box plots of Repli-seq signal for Group I and Group II genes across four S-phase fractions (S1–S4). Statistics by paired Wilcoxon test (A, B and C).

**Figure S6.**
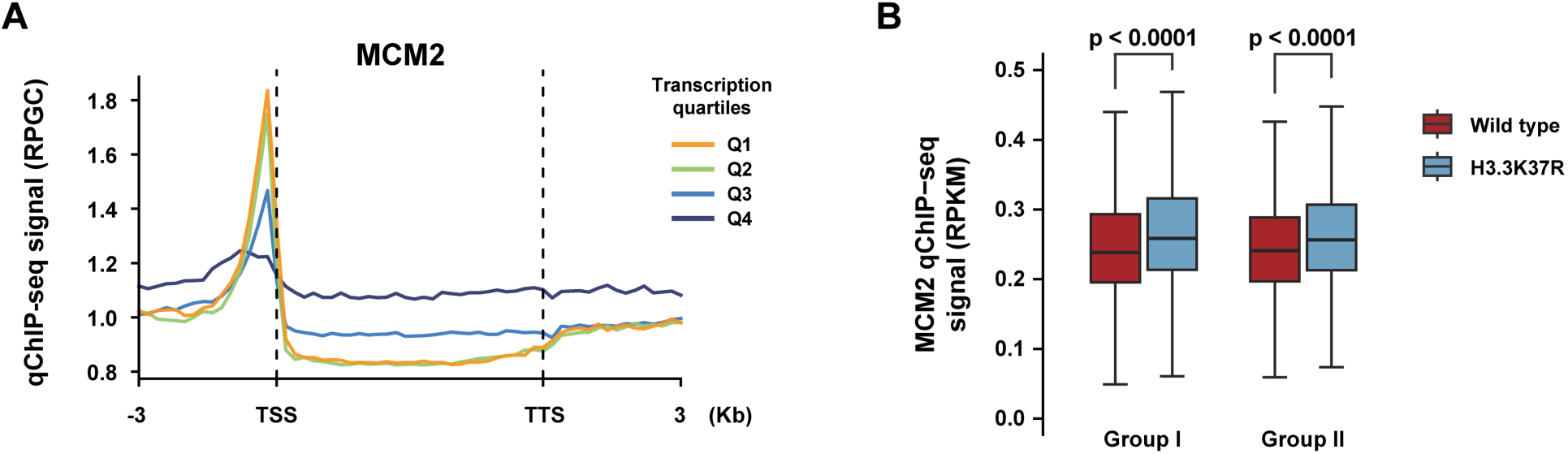
Supplementary to H3K37me1 limits MCM association with transcribed genes. **A.** Metaplot of average wild-type MCM2 qChIP-seq signal across protein-coding genes larger than 30 kb grouped into quartiles (Q1–Q4) by transcription levels. **B.** Box plot of MCM2 qChIP-seq signal for Group I and Group II genes in wild-type and H3.3K37R mutant cells. Statistics by paired Wilcoxon test (B).

